# Towards sustainable bioplastic production in resource limited environments using the photoferroautotrophic and photoelectroautotrophic bacterium *Rhodopseudomonas palustris* TIE-1

**DOI:** 10.1101/214551

**Authors:** Tahina Onina Ranaivoarisoa, Karthikeyan Rengasamy, Michael S. Guzman, Rajesh Singh, Arpita Bose

## Abstract

Bioplastics are an attractive alternative to petroleum-derived plastics because of the harmful environmental effects of conventional plastics and the impending fossil fuel crisis. Polyhydroxybutyrate (PHB) is a well-known bioplastic that is produced by several microbes using organic carbon sources. Autotrophic (using carbon dioxide or CO_2_) PHB production is reported for only a few organisms. Sustainable PHB bioproduction using other autotrophic microbes needs to be explored. *Rhodopseudomonas palustris*, a metabolically versatile purple non-sulfur bacterium (PNSB) is known to produce PHBs under photoheterotrophic conditions. *Rhodopseudomonas palustris* strain TIE-1 demonstrates extended metabolic versatility by using electron sources such as ferrous iron and poised electrodes for photoautotrophy. Here we report the ability of TIE-1 to produce PHB under photoferroautotrophic (light - energy source, ferrous iron - electron source and CO_2_ - carbon source) and photoelectroautotrophic (light - energy source, poised electrodes - electron source and CO_2_ - carbon source) growth conditions. PHB accumulation was observed both under nitrogen (N_2_) fixing and non-N_2_ fixing conditions. For comparison, we determined PHB production under chemoheterotrophic, photoheterotrophic and photoautotrophic conditions using hydrogen as the electron donor. Photoferroautotrophic and photoelectroautotrophic PHB production was on par with that observed from organic carbon substrates such as butyrate. PHB production increased during N_2_ fixation under photoheterotrophic conditions but not during photoautotrophic growth. Electron microscopy confirmed that TIE-1 cells accumulate PHBs internally under the conditions that showed highest production. However, gene expression analysis suggests that PHB cycle genes are not differentially regulated despite observable changes in biopolymer production.

## IMPORTANCE

PHB bioproduction was reported nearly a century ago. Despite its remarkable properties, PHB’s market competitiveness is affected by high production costs. Use of waste products such as molasses and industrial food waste for microbial PHB production can lower costs. An alternative cost-effective strategy is to employ microbes that use abundant and renewable resources. Toward that end, we report the ability of *Rhodopseudomonas palustris* TIE-1 to produce PHB under various growth conditions including photoferroautotrophy and photoelectroautotrophy. Because of the abundance of iron, CO_2_ and light on Earth, photoferroautotrophy is a sustainable, carbon neutral and low-cost strategy for PHB production. Photoelectroautotrophic PHB production can also be a useful approach in areas that produce electricity sustainably. Overall, our observations open new doors for sustainable bioplastic production not only in resource-limited environments on Earth, but also during space exploration and for *in situ* resource utilization (ISRU) on other planets.

Polyhydroxybutyrate (PHB), a member of the polyhydroxyalkanoate (PHA) family is the most common and well-studied biopolymer produced by bacteria (1). PHB is a potent substitute for conventional petroleum-based plastics because of many desirable properties. These include thermoresistance, moldability and biodegradability (2). PHB is also useful in many medical applications such as drug delivery, reconstructive surgery and bone tissue scaffolding (3). However, PHB production is currently not cost-effective (4). In order to reduce production costs, researchers have used many different carbon sources, which fall into two categories: 1) food wastes including sugar beet, soy and palm oil molasses or 2) cheap pure substrates as feedstock for PHB bioproduction. In some cases, less cost-effective substrates such as glucose that compete as food sources have also been tested for PHB production (1).

Autotrophic PHB production has been demonstrated by only a handful of organisms and remains an underexplored strategy for sustainable and carbon neutral bioplastic production. As carbon dioxide (CO_2_) concentrations rise in our atmosphere, such bioproduction strategies need to be explored further. The chemoautotrophs that produce PHBs include *Ideonella* sp., and *Cupriavidus necator* (*Ralstonia eutropha*) (5). Photoautotrophs represent an even more attractive group of organisms for PHB bioproduction due to their ability to use solar energy for biosynthesis. To this end, researchers have shown that cyanobacteria can produce PHB while performing oxygenic photoautotrophy (6). Anoxygenic phototrophic bacteria expand the repertoire of electron donors that can be used for such bioproduction. Several research groups have reported that PNSB can produce PHBs during photoheterotrophic growth (7, 8, 9). *Rhodopseudomonas palustris* is a biotechnologically important PNSB that can produce hydrogen under various growth conditions. This ability has sparked research on *R. palustris* strains to understand how PHB biosynthesis influences biohydrogen production (10, 11). To the best of our knowledge, PHB bioproduction under photoautotrophic conditions using inorganic electron donors has not been explored systematically in PNSB (5, 12).

To fill this knowledge gap, here we investigated the ability of the photoautotrophic PNSB *Rhodopseudomonas palustris* TIE-1 to accumulate PHB. We chose TIE-1 because it demonstrates extraordinary metabolic versatility even when compared to other *R. palustris* strains. For instance, similar to other *R. palustris* strains TIE-1 can grow chemoheterotrophically in rich medium and photoheterotrophically using many different organic carbon sources (13). It can use many different inorganic electron donors for photoautotrophic growth; some inorganic electron donors such as hydrogen and thiosulfate are similar to those also used by other *R. palustris* strains; other inorganic electron donors such as ferrous iron and poised electrodes are unique to TIE-1 (13, 14). Like other *palustris* strains, TIE-1 can also fix N_2_ gas (13). Importantly, TIE-1 is the only genetically tractable photoferroautotroph and photoelectroautotroph available (13, 14). Together, these abilities make TIE-1 an ideal candidate for testing photoautotrophic PHB production using unique inorganic electron donors.

To investigate the production of PHB by TIE-1, the strain was grown under various conditions. These include aerobic chemoheterotrophic growth on yeast extract-peptone (YP); and photoheterotrophic growth (anaerobic) using different carbon sources under both non-nitrogen (N_2_) and N_2_-fixing conditions. Aerobic growth on YP had the longest generation time compared to photoheterotrophic growth (Supplementary Table S3). Interestingly, YP grown cells produced the highest amount of PHB (13.94 g/L) amongst all non-N_2_ fixing heterotrophic conditions tested (Supplementary Table S3 and Figure 1 Panel A (a and b). TIE-1 produced the lowest amount of PHB (1.76 g/L) on succinate under non-N_2_ fixing conditions. PHB production varied from 2.22 g/L to 3.19 g/L when cells were grown in acetate, butyrate and 3-hydroxybutyrate (Supplementary Table S3 and Figure 1 Panel Ab). These low PHB levels might be linked to the availability of fixed nitrogen and an abundant carbon source, which could cause TIE-1 to increase cell numbers but not accumulate PHB as a carbon reserve. We examined PHB production under N_2_ fixing conditions. In general, N_2_ fixation delayed cell growth and resulted in a longer lag phase (approximately 2 times higher) when compared to growth under non-N_2_ fixing conditions. We also observed an increase in the maximum optical density (OD_660_) under N_2_-fixing photoheterotrophic growth conditions with acetate and butyrate whereas no significant difference was observed with succinate and 3-hydroxybutyrate (Supplementary Table S3). PHB accumulation increased on all N_2_-fixing photoheterotrophic conditions tested. PHB production doubled when cells were grown on 3-hydroxybutyrate (8.02 g/L), tripled on succinate (3.66 g/L), quadrupled on acetate (8.34 g/L), and increased 10-fold (23.12 g/L) on butyrate (Supplementary Table S3 and Figure 1 Panel Ab).

**Figure 1.**
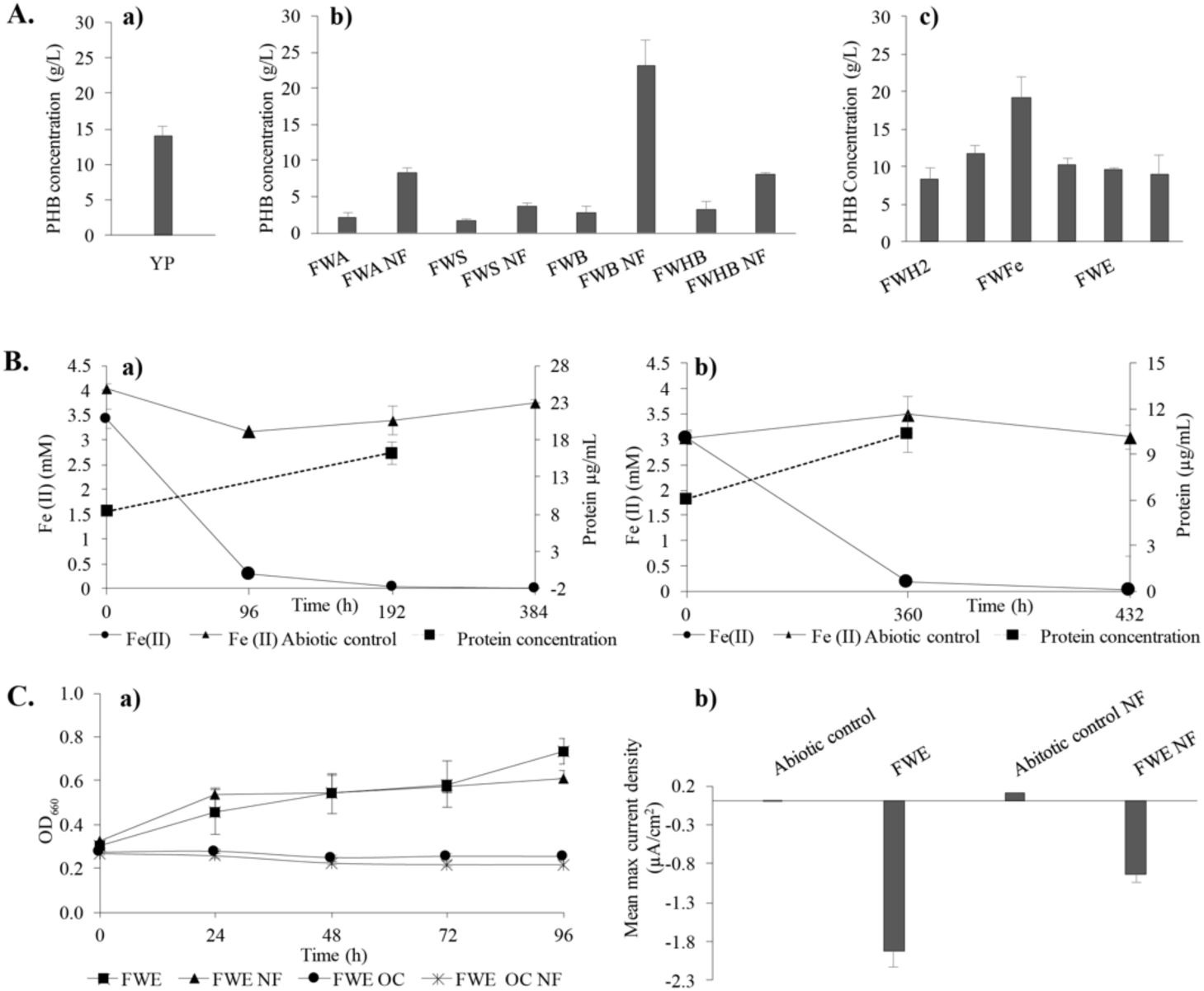
PHB production by TIE-1. **(a)** Under chemoheterotrophy in YP; **(b)** Under anaerobic photoheterotrophic conditions with ammonium chloride (NH_4_Cl) or under nitrogen fixing conditions (NF). Cells were grown with Fresh Water (FW) media supplemented with 10 mM succinate (FWS), acetate (FWA), butyrate (FWB) or hydroxybutyrate (FWHB). Cells were grown to late exponential phase: OD_660_~0.7; **(c)** Under anaerobic photoautotrophic conditions in H_2_CO_2_ (FWH2) grown to an of OD_660_~0.7, in 5 mM Iron (II) chloride (FWFe) for 8 days of growth with NH_4_Cl, 15 days of growth under nitrogen fixing condition (FWFe NF) and photoelectroautotrophy with NH_4_Cl (FWE), and for 4 days under nitrogen fixing conditions (FWE NF) (n=3 biological replicates). **Panel B.** Iron (II) oxidation and protein concentration during the growth of TIE-1 under photoferroautotrophic growth. Cells were grown in FW media supplemented with 5 mM Iron (II) chloride and NH_4_Cl in panel **(a);** and under nitrogen fixation condition in panel **(b)**. (n=3 biological replicates). **Panel C. (a)** OD_660_ values of the growth of TIE-1 grown under photoelectroautotrophy in FW supplemented with NH_4_Cl (FWE), under nitrogen fixing condition (FWE NF) or when the electrodes are not poised, called open circuit (OC). The two open circuit conditions shown are FWE OC and FWE OC NF; **(b)** Mean maximum current density (µA/cm^2^) (n=3 biological replicates) of FWE, FWE NF conditions, and the abiotic controls.

We also examined PHB production under photoautotrophic growth conditions using three different electron donors: hydrogen (H_2_), ferrous iron (photoferroautotrophy) and a poised graphite electrode (photoelectroautotrophy). TIE-1 was adapted to photoautotrophic growth using H_2_ in the presence or absence of fixed nitrogen. Under N_2_ fixing conditions, we observed that PHB production was as high as that observed during aerobic growth on YP (Figure 1 Panel Ac and Supplementary Table S3). Using H_2_, TIE-1 produced 11.69 g/L PHB under N_2_ fixing conditions and 8.4 g/L under non-N_2_ fixing conditions (Figure 1 Panel Ac and Supplementary Table S3). Growth on H_2_ under non-N_2_ fixing conditions was found to favor a higher maximal optical density (OD_660_ = 1.16) and a shorter generation time (*g* = 34 hours) when compared to N_2_ fixing conditions (OD_660_ = 0.54, *g* = 41 hours) (Supplementary Table S3). Photoferroautotrophic growth under non-N_2_ fixing conditions supported higher PHB production levels than those observed under YP growth, and close to those observed under the most productive photoheterotrophic condition (i.e. butyrate under non-N_2_ fixing conditions) (Figure 1 Panel Ac). The presence of PHB intracellular granules was confirmed by scanning transmission electron microscopy-electron energy loss spectroscopy (STEM-EELS) (Figure 2, Panel B). Under photoelectroautotrophic conditions the generation time did not change significantly under N_2_ fixing (*g* = 82 hours) vs non-N_2_ fixing conditions (*g* = 76 hours). TIE-1 grown in an open circuit reactor (electrode not passing current) did not show any growth under N_2_-fixing or non-N_2_ fixing growth conditions (Supplementary Table S3 and Figure 1 Panel Ca). After 96 hours of growth, the electron uptake was almost half under N_2_ fixing (0.93 µA/cm^2^) compared to non-N_2_ fixing conditions (1.92 µA/cm^2^) (Figure 1 Panel Cb and Supplementary Table S4). Slightly lower maximum planktonic OD_660_ was obtained under N_2_-fixing conditions compared to growth with fixed N_2._ The PHB concentration was 9.1 g/L under N_2_ fixing conditions and 9.6 g/L under non-N_2_ fixing conditions (Figure 1 Panel Ac, Panel Cb and Supplementary Table S3). When grown in an open circuit reactor, no PHB accumulation was detected (Supplementary Table S3) suggesting that PHB was used for cellular maintenance in the absence of an electron donor. In addition to carbon storage, PHB is suggested to act as an electron sink for bacteria especially under N_2_ fixing conditions (15). During photoferroautotrophic and photoelectroautotrophic growth, TIE-1 cells might be highly reduced. Under these conditions PHB biosynthesis could provide an electron sink, thus explaining the high level of PHB accumulation. McKinlay *et al.* have suggested a similar role for PHB synthesis in *R. palustris* CGA009 when it is grown under N_2_ deplete conditions with acetate (11).

**Figure 2.**
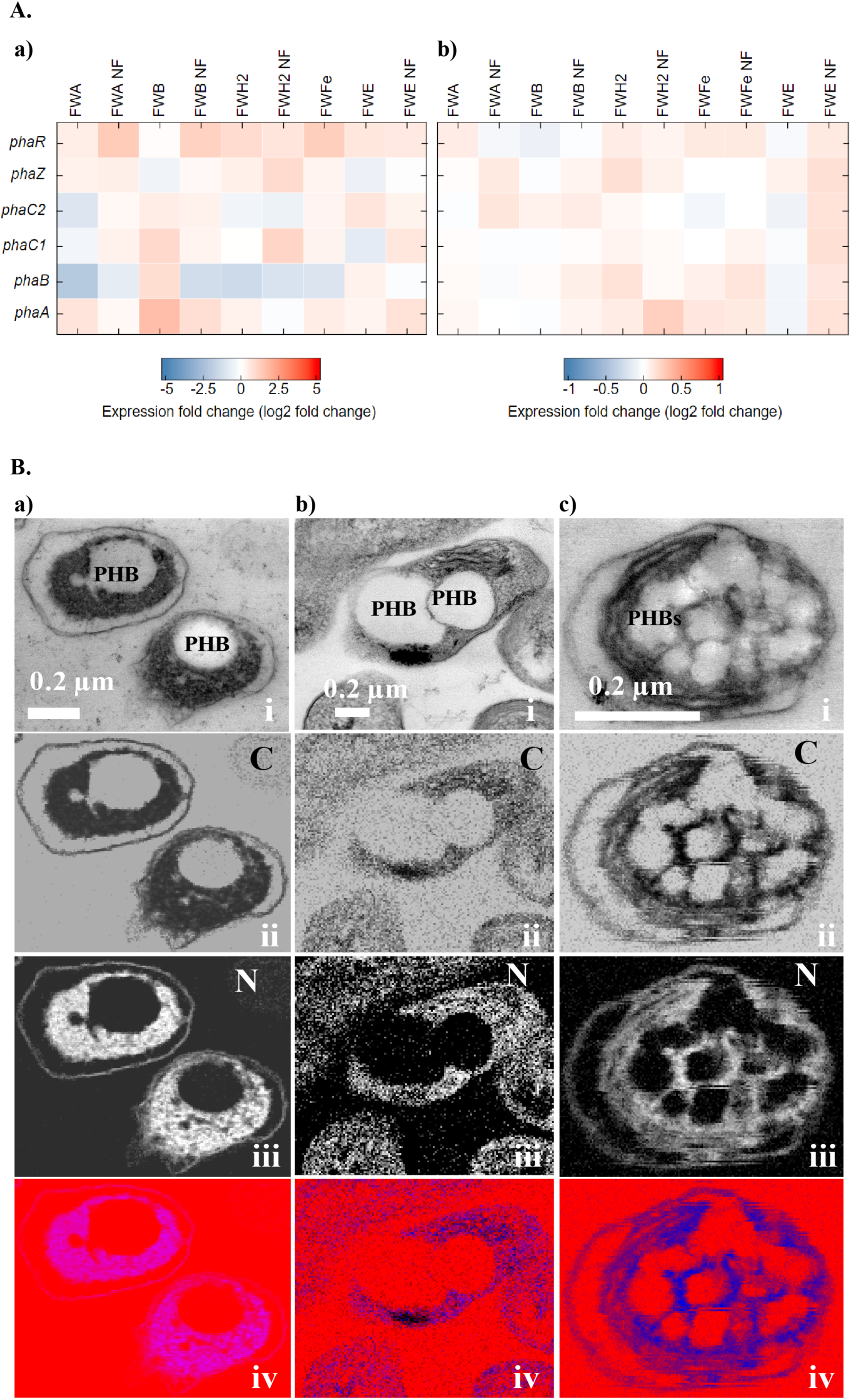
**Panel A.** Heat map showing log2 fold change in expression of PHB genes. **(a)** from RNA sequencing analysis; **(b)** from RT-qPCR. Results are from different growth conditions as described previously in Fresh Water (FW) media, either photoheterotrophically with succinate (FWS), acetate (FWA), butyrate (FWB), hydroxybutyrate (FWHB), or photoautotrophically with H_2_ (FWH2), photoferroautotrophically (FWFe), or photoelectroautotrophically (FWE) under N_2_-fixing (NF) or non-N_2_ fixing conditions. Results are from n=3 biological replicates. **Panel B.** STEM-EELS images of TIE-1 with PHB granules and corresponding carbon and nitrogen maps grown under FW media photoheterotrophically with butyrate in panel **(a)**; photoferroautotrophically in panel **(b)**; and photoelectroautotrophically in panel **(c)**. Bright areas represent the dominance of the corresponding element. Each panel comprises, (i) TEM bright field image, (ii) a nitrogen map (N) (iii) a carbon map (C) and (iv) a composite image where red represents carbon, blue nitrogen. The carbon background is due to the carbon-based resin that was used for embedding the cells.

To determine whether the expression of the genes involved in PHB biosynthesis and degradation change with changes in PHB levels, we performed transcriptomic analysis using RNA-Seq, and reverse transcription quantitative PCR (RT-qPCR). The genes include those encoding the first enzyme β-ketothiolase (acetyl-CoA acetyltransferase - PhaA that condenses two acetyl CoA molecules into acetoaetyl CoA; the second enzyme, an acetoacetyl CoA reductase that catalyzes the formation of the 3-hydroxybutyrate monomer from acetoacetyl CoA (PhaB); and finally, the polymerase (PhaC) that synthesizes the polymer PHB. When bacteria mobilize PHB, depolymerization of the granules is performed by the PHB depolymerase (PhaZ). The genes involved in the PHB biosynthesis pathway are well characterized in the PHB-producing model bacterium *Ralstonia eutropha* (16, 17, 18). The potential roles of the TIE-1 homologs are summarized in a simplified PHB cycle depicted in Supplementary Figure 1. Similar to *R. eutropha* (19), and *Bradyrhizobium japonicum* (20), there are multiple isozymes for the PhaA and PhaC enzymes, respectively, in TIE-1. In *Bradyrhizobium diazotoefficiens*, an organism closely related to *R. palustris*, PhaR regulates PHB biosynthesis by repressing the expression of *phaC*_*1*_ and *phaC*_*2.*_ In addition, PhaR regulates PhaP, a protein that binds to the surface and controls the number and size of the PHB granules. PhaR also binds to PHB granules during PHB synthesis and dissociates from it as the granule size grows (21). RNA-Seq analysis showed that the genes identified in the PHB cycle are not differentially regulated with respect to growth conditions or levels of PHB (Supplementary Table S5, S6 and S7). RT-qPCR analysis was performed on the *phaA* isozyme (Rpal_0532) that showed highest expression. Next to this *phaA* gene is a *phaB* isozyme gene with the locus tag Rpal_0533 (Supplemental Figure S1b, and Supplementary Table S8). The fact that these two genes appeared to be part of an operon supported our selection (Supplemental Figure S1b). Moreover, the gene for *phaR* (Rpal_0531) is next to the *phaA* (Rpal_0532) gene but expressed from the opposite strand. RT-qPCR corroborated the RNA-Seq data, which together show that the expression of the genes in the PHB cycle does not change under different growth conditions (Figure 2 Panel Aa-b). McKinlay *et al.* reported similar results when *R. palustris* CG009 was grown photoheterotrophically on acetate, under N_2_ deprivation (11). They observed that although PHB accumulated under these conditions, no change in PHB biosynthesis genes was notable (11).

## Implications

Here, we report PHB production by *R. palustris* TIE-1 under chemoheterotrophic (in YP); photoheterotrophic (using organic carbon sources: succinate, acetate, butyrate or 3-hydroxybutyrate); and photoautotrophic (using hydrogen, ferrous iron or poised electrodes as electron donor) conditions. Photoheterotrophic growth under N_2_ fixing conditions yielded higher PHB. The highest PHB production was obtained from photoheterotrophic growth on butyrate under N_2_ fixing conditions (23.12 g/L). This production is just 2-fold lower than the chemotrophic organism, *Cupriavidus necator* NCIMB 11599 (41.5 g/L) when it’s grown with a wheat based rich medium under nitrogen limited conditions (12). When compared to other autotrophs, the ability of TIE-1 to produce PHB under various heterotrophic, chemotrophic and phototrophic conditions offers a clear advantage in areas either depleted in organic carbon and/or having waste products as the most available carbon. In addition to its ability to produce PHB under heterotrophic conditions, TIE-1 can produce PHBs under unique photoautotrophic growth condition using H_2_, ferrous iron or poised electrodes as electron source. Furthermore, TIE-1 has the ability to fix atmospheric N_2_ which allows it to produce PHBs in the complete absence of available fixed nitrogen. We observed that anoxygenic photoautotrophic PHB production using abundant and accessible electron donors, such as ferrous iron, was higher than production under chemoheterotrophic growth in rich media.

Our results expand the substrate range that can be used by microbes for PHB production. TIE-1’s unique metabolic abilities such as photoferroautotrophy and photoelectroautotrophy can be used in novel sustainable PHB bioproduction platforms. Iron is the fourth most abundant element on Earth, and by using iron with solar energy, TIE-1 can be used to produce biochemicals such as PHB while capturing the potent and abundant greenhouse gas, CO_2_ (22, 23). Such a strategy will be especially valuable in resource-limited environments on Earth. Electricity can be produced renewably using solar and wind energy in many parts of our planet (24). In combination with solar energy and CO_2_, TIE-1 can be used to produce PHB using renewable electricity. Because TIE-1 can produce PHBs using many organic carbon sources, the use of waste materials as substrates is conceivable. Thus, TIE-1 represents a very metabolically versatile microbe that should be explored further for biochemical production not only on Earth but also during space exploration, and for *in situ* resource utilization (ISRU) on planets like Mars rich in iron, sunlight, carbon dioxide and nitrogen (25). TIE-1 based bioproduction strategies should be also be considered for waste management on Earth and “in flight” and “post arrival” during space exploration.

## Accession number(s)

All RNA sequence reads have been deposited with NCBI under BioProject accession number PRJNA417278.

## Acknowledgements

We are grateful to Josh Blodgett and Yifei Hu for their support during PHB analysis using LC-MS. We appreciate funding provided by The David and Lucile Packard Foundation as well as the Department of Energy (grant number DESC0014613).

## References

1. Verlinden RA, Hill DJ, Kenward MA, Williams CD, Radecka I. 2007. Bacterial synthesis of biodegradable polyhydroxyalkanoates. J. Appl. Microbiol. 102:1437–49.

2. Chen G-Q. 2010. Plastics from bacteria: natural functions and applications. Springer, Heidelberg; New York.

3. Manavitehrani I, Fathi A, Badr H, Daly S, Negahi Shirazi A, Dehghani F. 2016. Biomedical applications of biodegradable polyesters. Polymers 8:20.

4. Sabbagh F, Muhamad II. 2017. Production of poly-hydroxyalkanoate as secondary metabolite with main focus on sustainable energy. Renew. Sus. Energ. Rev. 72:95–104

5. Khosravi-Darani K, Mokhtari Z-B, Amai T, Tanaka K. 2013. Microbial production of poly(hydroxybutyrate) from C1 carbon sources. Appl. Microbiol. Biotechnol. 97:1407–1424

6. Troschl C, Meixner K, Drosg B. 2017. Cyanobacterial PHA Production-Review of Recent Advances and a Summary of Three Years' Working Experience Running a Pilot Plant. Bioeng. (Basel). 4(2):26

7. Mukhopadhyay M, Patra A, Paul AK. 2005. Production of poly(3-hydroxybutyrate) and poly(3-hydroxybutyrate-co-3-hydroxyvalerate) by *Rhodopseudomonas palustris* SP5212. World J. Microbiol. Biotechnol. 21(5):765–769

8. Mukhopadhyay M, Patra A, Paul AK. 2013. Phototrophic Growth and Accumulation of Poly(3-hydroxybutyrate-co-3-hydroxyvalerate) by Purple Nonsulfur Bacterium Rhodopseudomonas palustris SP5212. J. Polym. 2013:Aritical ID 523941.

9. Merugu R, Girisham S., Reddy S.M. 2010. Production of PHB (Polyhydroxybutyrate) by *Rhodopseudomonas palustris* KU003 under nitrogen limitation. Int. J. Appl. Biol. Pharm. Technol. 1(2) 676–678

10. Wu SC, Liou SZ, Lee CM. 2012. Correlation between bio-hydrogen production and polyhydroxybutyrate (PHB) synthesis by *Rhodopseudomonas palustris* WP3-5. Bioresour. Technol. 113:44–50

11. McKinlay JB, Oda Y, Ruhl M, Posto AL, Sauer U, Harwood CS. 2014. Non-growing *Rhodopseudomonas palustris* increases the hydrogen gas yield from acetate by shifting from the glyoxylate shunt to the tricarboxylic acid cycle. J. Biol. Chem. 289:1960–70.

12. Pagliano G, Ventorino V, Panico A, Pepe O. 2017. Integrated systems for biopolymers and bioenergy production from organic waste and by-products: a review of microbial processes. Biotechnol. Biofuel. 10:113.

13. Jiao Y, Kappler A, Croal LR, Newman DK. 2005. Isolation and characterization of a genetically tractable photoautotrophic Fe(II)-oxidizing bacterium, *Rhodopseudomonas palustris* strain TIE-1. Appl. Env. Microbiol. 71:4487–96.

14. Bose A, Gardel EJ, Vidoudez C, Parra EA, Girguis PR. 2014. Electron uptake by iron-oxidizing phototrophic bacteria. Nat. Commun. 5:3391.

15. Dawes EA. 1988. Polyhydroxybutyrate: an intriguing biopolymer. Biosci. Rep. 8:537–47.

16. Peoples OP, Sinskey AJ. 1989. Poly-beta-hydroxybutyrate biosynthesis in *Alcaligenes eutrophus* H16. Characterization of the genes encoding beta-ketothiolase and acetoacetyl-CoA reductase. J. Biol. Chem. 264:15293–7.

17. Peoples OP, Sinskey AJ. 1989. Poly-beta-hydroxybutyrate (PHB) biosynthesis in *Alcaligenes eutrophus* H16. Identification and characterization of the PHB polymerase gene (phbC). J. Biol. Chem. 264:15298–7.

18. Uchino K, Saito T, Jendrossek D. 2008. Poly(3-hydroxybutyrate) (PHB) depolymerase PhaZa1 is involved in mobilization of accumulated PHB in *Ralstonia eutropha* H16. Appl. Env. Microbiol. 74:1058–1063.

19. Slater S, Houmiel KL, Tran M, Mitsky TA, Taylor NB, Padgette SR, Gruys KJ. 1998. Multiple beta-ketothiolases mediate poly(beta-hydroxyalkanoate) copolymer synthesis in *Ralstonia eutropha*. J. Bacteriol. 180:1979–87.

20. Quelas JI, Mongiardini EJ, Perez-Gimenez J, Parisi G, Lodeiro AR. 2013. Analysis of Two Polyhydroxyalkanoate Synthases in *Bradyrhizobium japonicum* USDA 110. J. Bacteriol. 195:3145–3155.

21. Quelas JI, Mesa S, Mongiardini EJ, Jendrossek D, Lodeiro AR. 2016. Regulation of Polyhydroxybutyrate Synthesis in the Soil Bacterium *Bradyrhizobium diazoefficiens*. Appl. Env. Microbiol. 82:4299–4308.

22. Greenwood NN, Earnshaw A. 1984. Chemistry of the elements. Pergamon Press, Oxford.

23. Parida B, Iniyan S, Goic R. 2011. A review of solar photovoltaic technologies. Renew. Sus. Energ. Rev. 15:1625–1636

24. https://www.scientificamerican.com/article/u-s-reports-a-major-milestone-in-wind-and292solar-power/

25. Menezes AA, Cumbers J, Hogan JA, Arkin AP. 2015. Towards synthetic biological approaches to resource utilization on space missions. J. R. Soc. Interface 12:102.

